# Bayesian multiple logistic regression for case-control GWAS

**DOI:** 10.1101/198911

**Authors:** Saikat Banerjee, Lingyao Zeng, Heribert Schunkert, Johannes Söding

## Abstract

Genetic variants in genome-wide association studies (GWAS) are tested for disease association mostly using simple regression, one variant at a time. Standard approaches to improve power in detecting disease-associated SNPs use multiple regression with Bayesian variable selection in which a sparsity-enforcing prior on effect sizes is used to avoid overtraining and all effect sizes are integrated out for posterior inference. For binary traits, the *logistic* model has not yielded clear improvements over the linear model. For multi-SNP analysis, the logistic model required costly and technically challenging MCMC sampling to perform the integration.

Here, we introduce the *quasi-Laplace* approximation to solve the integral and avoid MCMC sampling. We expect the logistic model to perform much better than multiple linear regression except when predicted disease risks are spread closely around 0.5, because only close to its inflection point can the logistic function be well approximated by a linear function. Indeed, in extensive benchmarks with simulated phenotypes and real genotypes, our Bayesian multiple LOgistic REgression method (B-LORE) showed considerable improvements (1) when regressing on many variants in multiple loci at heritabilities ≥ 0.4 and (2) for unbalanced case-control ratios.

B-LORE also enables meta-analysis by approximating the likelihood functions of individual studies by multivariate normal distributions, using their means and covariance matrices as summary statistics. Our work should make sparse multiple logistic regression attractive also for other applications with binary target variables. B-LORE is freely available from: https://github.com/soedinglab/b-lore.

## Introduction

Common, noninfectious diseases are responsible for over ^2^/3 of the deaths worldwide. Genome wide association studies (GWAS) have opened up a fundamentally new approach to identify novel regions of the genome which are associated with these complex human diseases. In the past decade, GWAS identified thousands of genetic variants, particularly single nucleotide polymorphisms (SNPs), associated with many diseases and complex traits [1,2].

In a typical GWAS, genotype data comprising millions of SNPs from thousands of individuals with some trait are analyzed to identify SNPs that have significant associations with the trait. Most studies apply simple regression (single-SNP analysis) *i.e.*, they test for one SNP at a time, yielding a *p*-value for each SNP. GWAS for quantitative traits like lipid levels, BMI, height, etc. use a linear model for regression of the trait by the minor allele frequency at the SNP. Case-control GWAS, for which the binary trait is either “diseased” (“cases”) or “healthy” (“controls”), use a logistic model for regression.

Complex diseases studied with GWAS are usually polygenic, with many SNPs each contributing only a small fraction of the disease risk. The low effect sizes limit the power to detect statistically significant associations. Meta-analyses attempt to increase the statistical power by combining the summary statistics from single-SNP analyses of many GWAS, thereby encompassing a large number of samples, often in the range of hundreds of thousands.

While the simple regression model is computationally fast, it can only detect association and not statistical coupling or even causality. Therefore, a non-causal SNP (tag SNP) that is instrong linkage disequilibrium (LD) with a causal SNP will also obtain similarly significant *p*-values, making it difficult to decide which of these SNPs is really causal. Multiple regression models (also called multiple-SNP analyses or polygenic models) overcome this problem by using many SNPs at a genetic region or locus as explanatory variables. They can distinguish between correlation and coupling because the causal SNP can explain away the effects of other SNPs, which are merely correlated with the given phenotype via the causal SNP. Multiple regression can also improve the power of GWAS to detect risk loci by aggregating evidence from many SNPs with low effect sizes.

In GWAS, only a few hundred out of millions of measured or imputed SNPs are expected to be causal, *i.e.*, to have a direct influence on disease risk. Therefore, Bayesian variable selection has been employed to prevent overtraining (see [3] for an overview). Bayesian variable selection uses a sparsity-enforcing prior on the effect sizes to force all but a small fraction of the regression coefficients to zero. Bayesian variable selection regression (BVSR) [4,5] uses a point-normal prior, a mixture of a delta function at zero and a normal distribution for causal SNPs.

Until recently, multiple regression was limited by the requirement of individual-level genotype data. It was practically infeasible to apply it to multiple GWAS due to logistical, technical, and ethical restrictions for sharing huge volumes of genetic data from patients. In recent years, a number of studies devised novel strategies to perform multiple regression with variable selection using only summary statistics: PAINTOR [6,7], CAVIAR [8], CAVIARBF [9] and FINEMAP [10] employ a linear model yielding a multivariate normal likelihood. Its mean is approximated using the single-SNP effect size estimates and its covariance matrix is deduced from the LD correlation matrix of the SNPs. Like BVSR, they use a point-normal prior for variable selection. These methods are routinely used for fine-mapping i.e., for prioritizing the SNPs within the risk-associated loci (see [11] for a recent review).

For binary phenotypes, these fine-mapping methods approximate the logistic likelihood with a linear function of the genotype vector, permitting an analytical solution of the integral over effect sizes. Because the integration is analytically intractable without this linear approximation, multiple *logistic* regression required computationally cumbersome Markov Chain Monte Carlo (MCMC) sampling. Early in 2009, Newcombe et *al.* developed a Bayesian framework for multiple logistic regression using variable selection [12] using full MCMC sampling of all parameters and analyzing ~35 SNPs. For analyzing binary traits with BVSR [5], Guan and Stephens used the probit model, which is very similar to the logistic model. However, they could not demonstrate a clear benefit of the probit model and ascribed this to technical difficulties in the MCMC sampling (insufficient mixing for binary traits), concluding that the approach needs “further methodological innovation”.

Besides the methodological challenge of performing the integration over effect sizes, single-SNP logistic regression did not yield clear advantages over linear regression [13], which might have also reduced the interest of exploring multiple logistic regression. Here we argue that the reason for the lack of improvement is simply due to the fact that the risk explained by a single SNP is usually so low that predicted risks stay very near to 0.5, where a linear approximation of the logistic function is still very accurate.

In this work, we present B-LORE, a scalable Bayesian method for multiple logistic regression. We introduce the quasi-Laplace approximation in which we approximate the *L*_2_-regularized likelihood of the logistic model by a normal distribution, whose mean vector and covariance matrix serve as our novel summary statistics. This trick allows us to analytically integrate the (unregularized) likelihood times the point-normal effect-size prior over the unknown effect sizes. The regularization ensures that the mode of the regularized likelihood is near the mode of the integrand and hence the normal approximation stays accurate. We estimate the parameters of our effect-size prior by maximizing the total marginal likelihood over all loci. B-LORE can also combine multiple case-control GWAS because the maximization requires only the summary statistics for each study.

Through extensive benchmarks, in which we simulate binary phenotypes for real genotype data, we show that the quasi-Laplace approximation is significantly better than existing linear approximations of the logistic model. Most other multiple regression methods using Bayesian variable selection have been developed for fine-mapping of SNPs within credible sets, *i.e.*, sets of SNPs known to contain at least one causal SNP. We therefore limit our comparison here tothis application. However, B-LORE can also be employed to rank risk loci by their probability to contain a causal SNP or to predict the genetic risk of patients from their genotype.

## Materials and Methods

We are interested in analyzing case-control GWAS, using summary data instead of individual genotype data. In this section, we describe the model and the implementation of B-LORE. At each step we compare and contrast B-LORE with other multiple regression variable selection methods, namely BIMBAM [4], piMASS [5], GEMMA [3,14], CAVIARBF [9], FINEMAP [10] and PAINTOR [6,7]. For quick reference, we summarized the methods in Table 1. Finally, we describe the data and simulation details used for the validation of our method.

**Table 1.** Comparison of methods for multiple regression of case-control GWAS data. All methods use the point-normal prior (Eq. (4)) for the effect sizes of the SNPs. The posterior probability is obtained by integrating out these effect sizes, either via MCMC sampling or analytically (column 2). Column 3 compares the approximations to the logistic or probit likelihood model. Column 4 shows which hyperparameters of the prior distribution are estimated from the data, and the method of estimation is shown in column 5. Column 6 lists which tools can perform meta-analysis. MCMC: Markov Chain Monte Carlo sampling, EM: expectation-maximization, CG: conjugate gradient method.

| Method | Integration method | Approximation | Hyperparameters learnt from data | ... using method | Multiple loci | Meta-analysis |
| --- | --- | --- | --- | --- | --- | --- |
| BIMBAM [4] | MCMC | Laplace | None | – | No | No |
| piMASS [5] | MCMC | Probit | $\pi_i, \sigma$ | MCMC | No | No |
| GEMMA [3, 14] | MCMC | Probit | $\pi_i, \sigma$ | MCMC | No | No |
| CAVIARBF [9] | Analytic | Linear | None | – | No | Yes |
| FINEMAP [10] | Analytic | Linear | None | – | No | Yes |
| PAINTOR [6, 7] | Analytic | Linear | $\pi_i$ | EM | No | Yes |
| **B-LORE** | Analytic | quasi-Laplace | $\pi_i, \sigma$ | CG | Yes | Yes |

### Likelihood function

For binary traits, GWAS data consists of phenotypes *φ_n_* ∈ {0,1} (healthy or diseased) and of genotypes *w_ni_* ∈ {0,1, 2}, where 0, 1, or 2 signify the number of minor alleles of patient *n* ∈ {1,…, *N*} at SNP *i* ∈ {1,…, *I*}. The genotype is centered and normalized as 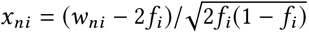, where *f_i_* is the minor allele frequency of the *i*^th^ SNP. We denote the vector of normalized genotypes for the *n*^th^ sample as *x_n_*, and the *N × I* matrix of genotypes as **X**.

As BIMBAM, CAVIARBF, FINEMAP and PAINTOR, we use standard logistic regression to model the probability for a patient to have the disease,

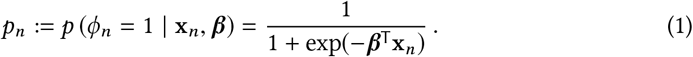

which can be transformed to

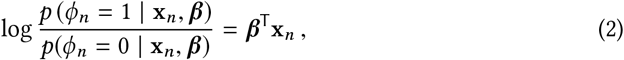

The minor allele count *x_ni_* of SNP *i* contributes linearly to the log-odds ratio with an effect size *β_i_*. The likelihood for *N* patients is

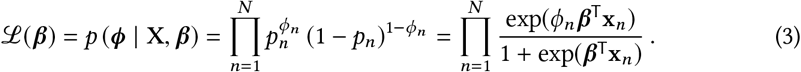

### Prior distributions

#### Point-normal prior for effect sizes

In a GWAS, the number of parameters *p = I* is usually much larger than the number of samples *N* (*p ≫ N*). Hence, a standard approach of maximizing the likelihood with respect to the effect sizes will lead to gross overtraining. One common solution is to add a regularization term to the log likelihood that will push most of the components of ***β*** to zero or near zero. From a Bayesian viewpoint, this is equivalent to maximizing the posterior distribution *p*(***β***|**X**), which is proportional to *p*(**X**|***β***) *p*(***β***), where *p*(***β***) is the prior distribution of effect sizes [15].

To reflect the prior expectation that an overwhelming majority of SNPs have a negligible effect on disease risk, we use the point-normal prior,

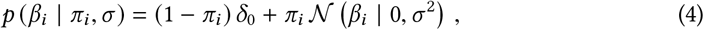

which is used in many tools, *e.g.* BIMBAM, piMASS, CAVIARBF, FINEMAP and PAINTOR. The normal distribution models the effect sizes for the rare, causal SNPs and the delta function models the non-causal SNPs. The hyperparameters *π_i_* control the sparsity of the model and *σ*^2^ describes the variance of the effect sizes.

#### Prior probability of *π* and *σ*

In the simplest case the prior probabilities to be causal is the same for all SNPs, *π_i_ = π* = const. CAVIARBF and FINEMAP implicitly assumes that *π_i_* = 1/I. B-LORE also assumes *p*(*π*) = const. However, improvements can be expected by making *π_i_* depend on informative local genomic features or annotation tracks [7].

BIMBAM, PAINTOR, CAVIARBF and FINEMAP use fixed values of *σ* that can be specified by the user. PiMASS uses an intuitively appealing prior on *σ* with wider tails (see [5] for details). B-LORE implicitly assumes a much simpler prior *p*(*σ*^2^) = const to avoid computational complexity.

#### Causality configurations

We define *c_i_* ∈ {0,1} as the hidden indicator variables defining the underlying causality of the SNPs. Here, *c_i_* = 1 indicates that SNP i is causal and *c_i_* = 0 otherwise. To simplify notations, we define the vector 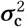, whose *i*^th^ component is 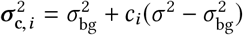. This allows us to reformulate Eq. (4) as:

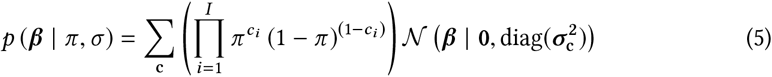

with the sum running over all 2^*I*^ possible *causality configurations* c ∈ {0, 1}^*I*^. Using

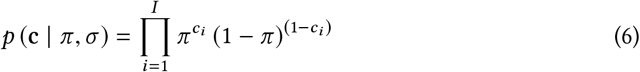

we can write the prior on the effect sizes as,

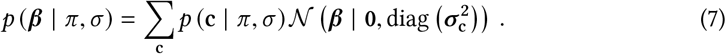

In this formulation, *p*(*c_i_* = 1 | *π, σ*) gives the prior probability of the *i*^th^ SNP to be causal before observing the phenotype and genotype data, and ║**c**║_1_ gives the total number of causal SNPs in the model.

BVSR and CAVIARBF also use the same Bernoulli prior on c (Eq. (6)). FINEMAP uses a general discrete distribution for the number of causal SNPs. However, it requires that the region to be analyzed includes at least one causal SNP, *i.e.*, *p* (c = 0) = 0. For binary traits, piMASS also has the same restriction.

### Inference

#### Fine-mapping

The posterior probability for SNP *i* to be coupled to the disease is obtained by summing the posterior probability over all causality configurations c for which SNP *i* is causal (*i.e., c_i_* = 1):

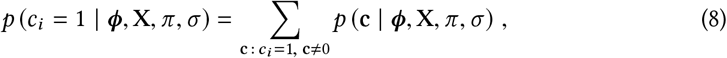

also called the posterior inclusion probability (PIP). CAVIARBF, FINEMAP and BVSR also outputs the PIP.

#### Prediction of causal loci

The probability for a locus to be coupled with the disease phenotype is equal to the probability of the locus harboring at least one causally associated SNP. This is equal to 1 minus the probability of not containing a single causal SNP:

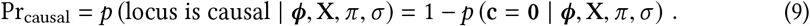

CAVIARBF and FINEMAP output Bayes factor for the posterior probability that there is at least one causal variant in the region against the null model.

### Quasi-Laplace approximation

Both the above posterior inferences require computing

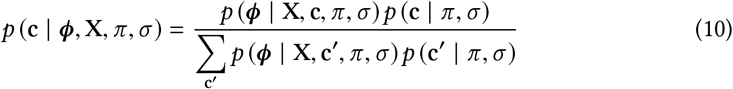

for all causality configurations c, which in turn requires computing

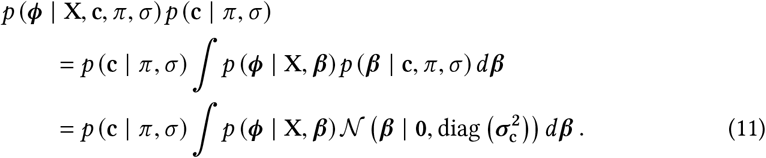

The above integration also appears in the marginal likelihood (see below) used for the optimization of the hyperparameters (*π, σ*). It does not have an exact solution when using the likelihood function of the logistic model, given by Eq. (3). In contrast to logistic regression, the linear regression has normally distributed likelihood function, admitting an exact solution (for example, see Protocol S1 of [4]).

BIMBAM approximates the integrand with a multivariate Gaussian using Laplace’s method. In this method, the parameters of the Gaussian are determined by finding the integrand’s mode (e.g. using gradient-based optimization) and setting the precision matrix to the Hessian at the mode. Unfortunately, the mode depends on c and (*π, σ*). So, one needs to determine the mode and precision matrix every time (*π, σ*) is changed. Not only does this require individual genotype data, but this also makes it computationally infeasible to learn (*π, σ*).

CAVIARBF and FINEMAP approximate *p_n_* with a linear function of ***β***, which reduces the likelihood function of Eq. (3) to a multivariate normal distribution with scaled variance (see Pirinen *et al.* [16] and Chen *et al.* [9] for details). This approximation becomes inaccurate as we move away from the mode of the likelihood, and unfortunately the region in ***β*** space which contributes most to the integrand (around the mode of the integrand) can be quite far from the mode of the likelihood.

We propose the “quasi-Laplace approximation” to solve the integration. We start by splitting the integrand into two factors – an *L_2_-regularized likelihood* that approximates the integrand (but does not depend on (*π, σ*) or c) and a correction term:

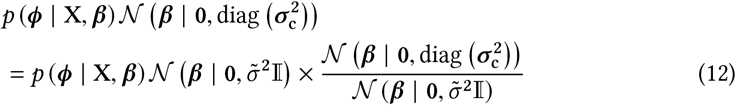

The regularized likelihood is the product of the likelihood function (Eq. (3)) and a Gaussian *regularizer*, 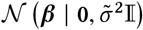 which acts as an approximate, simple prior distribution - pulling the maximum of the regularized likelihood near to the mode of the integral, making it more accurate than the Laplace approximation. The 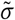 can be optimized on the data (see S1 File for details) or can be specified by the user. Next, we approximate the regularized likelihood by a multivariate Gaussian:

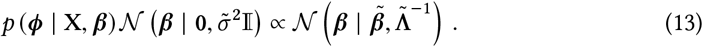

The quasi-Laplace approximation can be motivated from its validity in the limit of *N* ≫ 1 and equal number of cases and controls (details in S1 File for interested readers). From our simulations, we found that the approximation can be extended to unequal number of cases and controls. The regularized likelihood does not depend on (*π, σ*) or c. Hence, it would suffice to calculate it only once and use 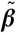 and 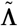 as our summary statistics. We determine 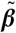 and 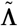 by maximizing the regularized likelihood with respect to ***β***, and setting 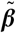 to the mode and 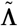 to the negative of the Hessian matrix at the mode. The covariance 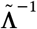 resembles the scaled LD matrix used for the logistic model in CAVIARBF, FINEMAP and PAINTOR, but includes an additional term from the regularizer (see S1 File).

### Optimization of hyperparameters

In the same spirit as BVSR, CAVIARBF and FINEMAP, we calculate the *marginal likelihood function* [15],

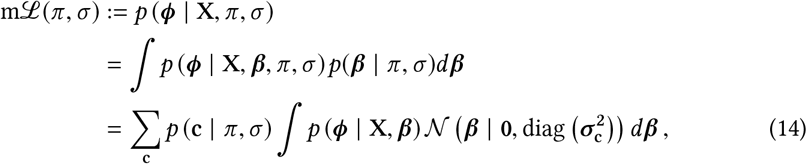

where we use Eq. (7) in the last step. In contrast to the classical maximum likelihood approach, in which the parameters ***β*** are optimized, this method integrates out the parameters ***β***. This is a crucial difference in practice, because it eliminates the need to learn a large number of parameters and thereby very effectively guards against overtraining. Also, by integrating out the parameters we avoid the errors incurred when fixing them to noisy point estimates.

In B-LORE, the integration in Eq. (14) is solved by the quasi-Laplace approximation whereas other methods use a linear approximation of logistic model to solve the integration.

The solution (see S1 File) depends on 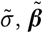 and 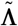, which serve as our novel summary statistics.

In B-LORE, we use the conjugate gradient method to maximize the marginal likelihood function. BIMBAM, CAVIARBF and FINEMAP use fixed values of the hyperparameters (*π, σ*). PAINTOR learns *π* from the data. PiMASS and GEMMA learn both *π* and *σ* from the data using MCMC.

### Meta-analysis

Meta-analysis would require combining 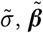 and 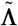 from multiple studies. For a single study, the likelihood is given by Eq. (3). We can combine multiple independent studies simply by computing the total likelihood as the product of the likelihoods of each contributing study *s*:

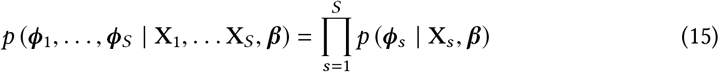

The integrand in Eq. (14) will now have a product over multiple logistic functions. We have the summary statistics 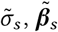 and 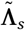 from each study. We apply the quasi-Laplace approximation for each study and combine the regularized likelihoods to a multivariate normal distribution:

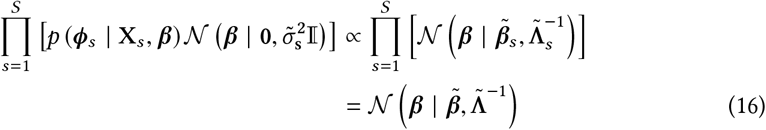

where we now have,

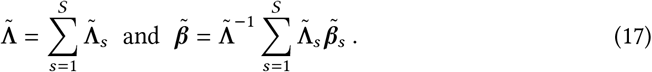

We now use Eq. (17) in place of Eq. (13) to calculate the marginal likelihood for the optimization of hyperparameters. Unlike conventional meta-analysis methods, which pool aggregate allele count data of each individual SNP, the above method allows us to combine information from multiple regression.

### Overview of steps used in B-LORE software

In summary, B-LORE works in two steps:

1. *Novel summary statistics.* Calculation of summary statistics in B-LORE requires two optimization at each cohort or study:

1. Learn 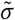 from the data.
2. Learn 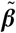 and 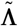 from the data.
3. *Meta-analysis.* Estimation of the hyperparameters (*π, σ*) by optimizing the marginal likelihood using 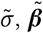 and 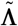 of each cohort or study.

In our software, the first step can be run using the command --summary and the second step can be run using the command --meta.

### Factorization over loci

To speed up B-LORE analysis, we recommend to pre-select loci with a faster method such as SNPTEST [17–19] and to include SNPs from these pre-selected loci. Usually these candidate loci will be in linkage equilibrium since LD is highly local. Therefore the covariance matrix X^⏉^X is approximately block-diagonal, with each block corresponding to a locus. All multiple regression methods utilize this block-diagonal feature of the LD matrix. For example, BIMBAM uses a factorization over loci to perform multiple regression at each locus independently.

However, analyzing multiple loci together increases power and specificity for fine-mapping [20]. We expect the logistic model to benefit from analyzing multiple loci because they can together explain a higher fraction of heritability. Hence, we compute the summary statistics over all the pre-selected loci together. In S1 File, we show that the marginal likelihood in Eq. (14) can be factorized as a product over all the loci. Therefore, the hyperparameter optimization can be performed over all loci simply by summing the log marginal likelihoods of all loci. This will effectively ignore the off-diagonal terms in the covariance matrix among the loci but retain the important diagonal elements.

### Data

We used the genotype from five German population cohorts: German Myocardial Infarction Family Study (GerMIFS) I - V [21–26]. Details for quality control and pre-processing of these data were described by Nikpay *et al.* [26]. Briefly, there are a total of 6234 cases and 6848 controls of white European ancestry. Each cohort was imputed with phased haplotypes from the 1000 Genomes Project. SNPs were filtered for MAF > 0.05 and Hardy-Weinberg equilibrium (HWE) *p*-value > 0.0001.

### Phenotype simulation

The inherent complexity of the genotype data with strong linkage effects are very difficult to simulate realistically from haplotype data. We therefore used real patient genotypes for our semi-synthetic benchmark. We selected 100 genomic regions or loci from each of the five GerMIFS cohorts, such that each locus had 200 unique SNPs.

Since piMASS cannot do meta-analysis, we had to make a *unified* cohort by combining the individual genotypes and phenotypes of the five cohorts. Hence, we pruned each locus to contain only SNPs which are common to all the five cohorts, leaving 17218 SNPs distributed over the 100 loci.

In each locus, we sampled one or more SNPs to be causal. S2 Figure shows the distribution of the number of causal SNPs at each locus. Once we established a “ground truth” of C causal SNPs, we used the classical liability threshold model [27, 28] to simulate the binary phenotypes *φ_n_* for every individual. Guan and Stephens [5] also used the same method for simulating binary phenotypes. The model assumes that the binary disease status results from an underlying continuous disease liability that is normally distributed in the population. If the combined effects of genetic and environmental influences push an individual’s liability across a certain threshold level, the individual is affected.

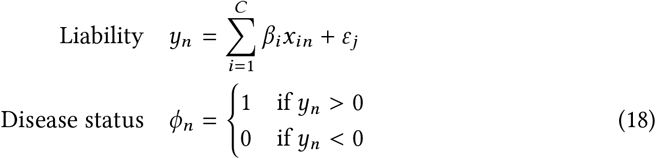

This is equivalent to a disease prevalence of 0.5 and gives roughly equal number of cases and controls. The individual effect sizes *β_i_* were assumed to be normally distributed 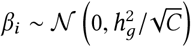 such that the causal SNPs aggregated to explain a fixed heritability (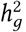, proportion of the phenotypic variance) on the liability scale.

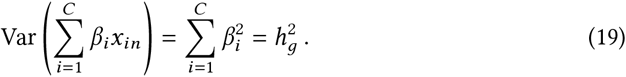

The environmental contribution given by *ϵ_j_*, was assumed to be normally distributed 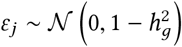. The observations on the risk scale follows a probit function of the liabilities on the unobserved continuous scale [28].

### Methods for comparison

Multiple regression methods are primarily used for fine-mapping, and are generally applied to credible sets, *i.e.*, loci which contain causal SNPs. Hence, we compared the ranking of SNPs within each locus. Fine-mapping methods are generally assessed in terms of recall, *i.e.*, the proportion of all causal SNPs in the locus included in the top ranked SNPs. In addition, we also looked at the precision, *i.e.*, the proportion of causal SNPs among the selected ones.

#### B-LORE

We calculated the summary statistics at each cohort and performed meta-analysis. B-LORE is not designed to be applied on a single locus. It learns the hyperparameters by conjugate gradients from the data and would require enough number of SNPs for proper estimation. A single locus generally do not have enough causal SNPs in real data. Hence, all loci were used for the meta-analysis in each simulation, unless otherwise stated. The SNPs were ranked by the PIPs within each individual locus.

#### META

We used META v1.7 for single-SNP meta-analysis. At each individual cohort, we obtained summary statistics with SNPTEST v2.5.2 assuming an additive model and using a missing data likelihood score test (`–method score`). We then corrected each cohort for the genomic inflation factor, and performed meta-analysis using inverse-variance method based on a fixed-effects model. We used the – log_10_ (*p*) values for ranking.

#### FINEMAP

We used FINEMAP v1.1 for multiple regression with a linear approximation to the logistic model. It has similar accuracy as CAVIARBF and higher accuracy than PAINTOR without functional annotations. As input, FINEMAP requires the *z*-score for every SNP and the LD matrix for each locus. We calculated 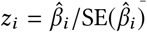 using the results of META. The prior standard deviation was set as 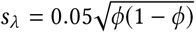, where φ is the proportion of cases among the *N* individuals. To ensure the best performance, we calculated the LD matrix using LDSTORE [29] from the single *unified* cohort. After analysis, we used the logarithm of the Bayes factors, log_10_ (BF (*c_i_* = 1: *c_i_* = 0)) for ranking the SNPs.

#### BVSR

GEMMA and piMASS both provide a BVSR probit model implementation using MCMC integration. The probit model is similar to the logistic model. We performed MCMC sampling at each locus of the *unified* cohort using piMASS with the `–cc` flag. We used 100000 burn-in steps and 1000000 production steps for the MCMC. We also repeated the same analysis without the `–cc` flag to compare the probit model with the linear model. We again used the PIPs for ranking the SNPs.

## Results

To evaluate the validity, accuracy and utility of the quasi-Laplace approximation, we performed a series of simulations to explore different conditions.

### Effect of heritability

First we show that B-LORE outperforms existing fine-mapping methods for binary phenotypes in case-control GWAS (Fig. 1). In general, the recall of every method improves with heritability. At all heritabilities, B-LORE provides the best ranking of SNPs followed by the multiple regression methods (BVSR and FINEMAP), which are better than single-SNP meta-analysis (META). The difference in recall between B-LORE and BVSR increases with increasing heritability – the improvement being always equal to or better than the difference between BVSR and META.

**Fig 1.**
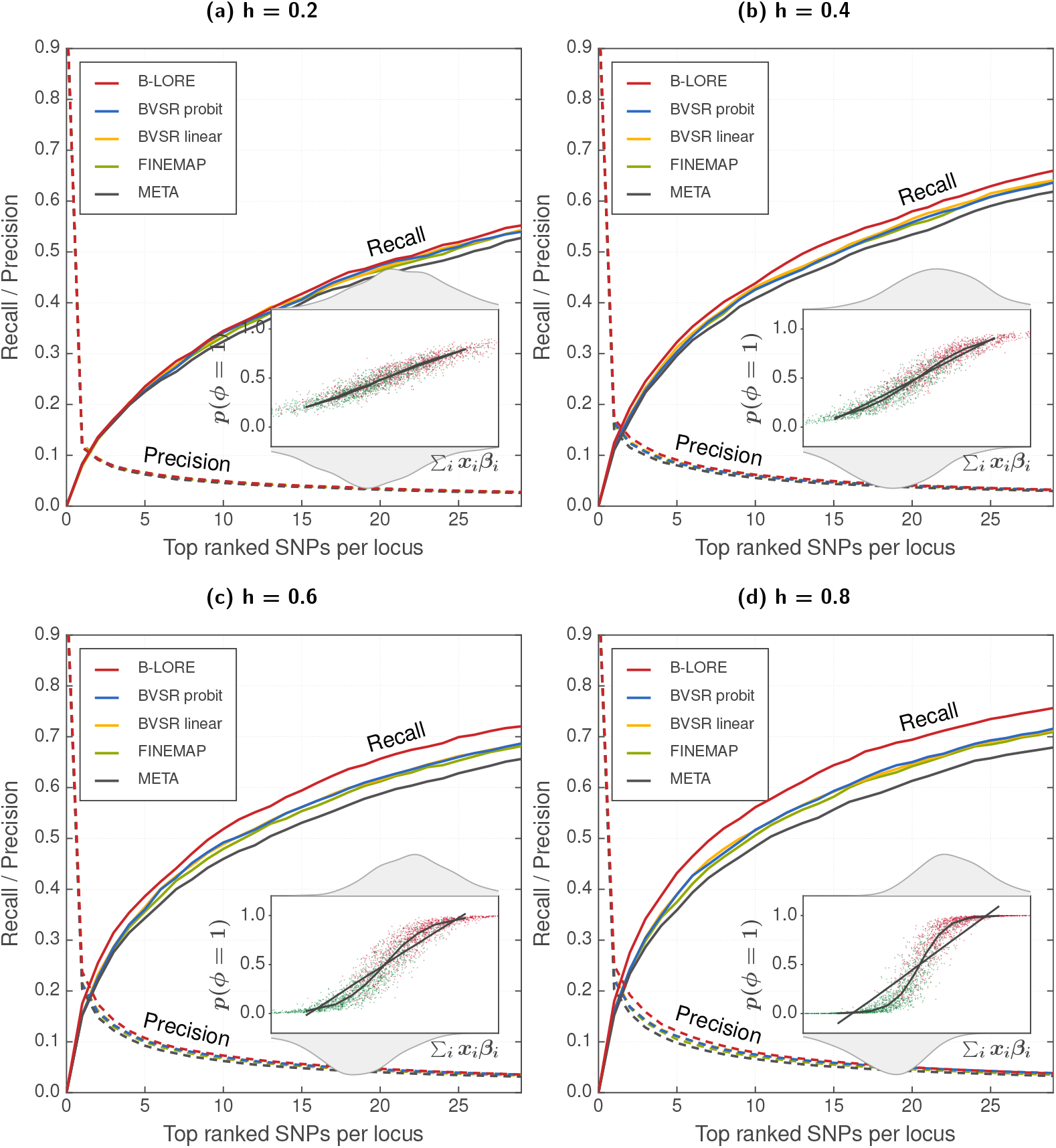
Multiple logistic regression improves fine-mapping in case-control GWAS. We simulated 13082 phenotypes using 100 loci of ~200 SNPs, as described in the main text. We compared the ranking of SNPs at each locus using recall (solid lines, left *y*-axis) and precision (dotted lines, right *y*-axis), which were averaged over 100 loci and 20 simulation replicates. All methods were run with a maximum of two causal SNPs per locus. Different panels show the results at different heritabilities, 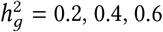 and 0.8. Insets schematically compare the logistic model with the linear model. We plot the true Σ_*i*_ *x_i_ β_i_* from the simulation for each individual along the *x*-axis, and show the distribution of cases and controls on the top and bottom axes respectively. On the *y*-axis, we show the predicted probability of being causal using a logistic model (*p* (*φ* = 1), red for cases and green for controls). The black lines are merely visual guides, obtained by averaging in each quantile. With increasing heritability, the predicted disease probability spreads away from 0.5, where the logistic model becomes increasingly better than the linear model to explain the data and B-LORE shows increasingly more recall over other methods. The improvement by B-LORE over other multi-SNP analyses is more significant than the improvement by multi-SNP over single-SNP analyses.

We schematically compare the logistic model against the linear model for our simulated GWAS (insets of Fig. 1). At low heritability, 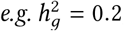, the cases and controls appear near the inflection point of the logistic curve, *i.e.*, in the linear regime, and hence the linear model is a good approximation of the logistic model. With increasing heritability, the cases and controls have a wider spread and increasingly more samples appear in the non-linear regime. Hence, the linear model becomes increasingly inaccurate for explaining the data. Therefore, the quasi-Laplace approximation performs better than the linear approximation of the logistic model.

B-LORE also outperforms the BVSR probit model. We kept B-LORE as similar as possible to the BVSR probit model in order to validate the quasi-Laplace approximation – using the same priors for ***β*** and *c* and a simpler hyperprior for *σ*. There are two major differences in the implementation. First, piMASS uses MCMC for optimization while B-LORE uses conjugate gradient method including the quasi-Laplace approximation. Second, piMASS analyzes each locus independently while B-LORE uses multiple loci. Therefore, the poorer performance of piMASS compared to B-LORE could be due to: (1) inefficient MCMC sampling by piMASS for binary phenotypes (as postulated by the authors), or (2) less information in each individual locus, as compared to the total heritability accessible to B-LORE from multiple loci (as postulated by Newcombe *et al.* [20] in a different context).

In the following we try to explore the contribution of the logistic model and of multiple loci for the improved performance of B-LORE.

### Effect of case-control imbalance

We reasoned that if the improved performance of B-LORE is due to the more accurate modelling of risk in the non-linear regime, then B-LORE should also excel when the bias constant is far from zero, *i.e.*, for case/control ratios less than 1.0. Having more controls than cases would lead to extreme imbalance in the data, making the linear approximation grossly suboptimal.

Our simulation yields roughly equal number of cases and controls – approximately ~6500 cases and ~6500 controls with a disease prevalence of 0.5. To obtain unequal cases and controls, we picked a subset of cases and/or controls by random choice. This can be done in two ways: (a) using the same number of cases (Fig. 2), and (b) using the same number of controls (S4 Figure). The remaining cases and/or controls were ignored by assigning unknown (NA) disease status.

**Fig 2.**
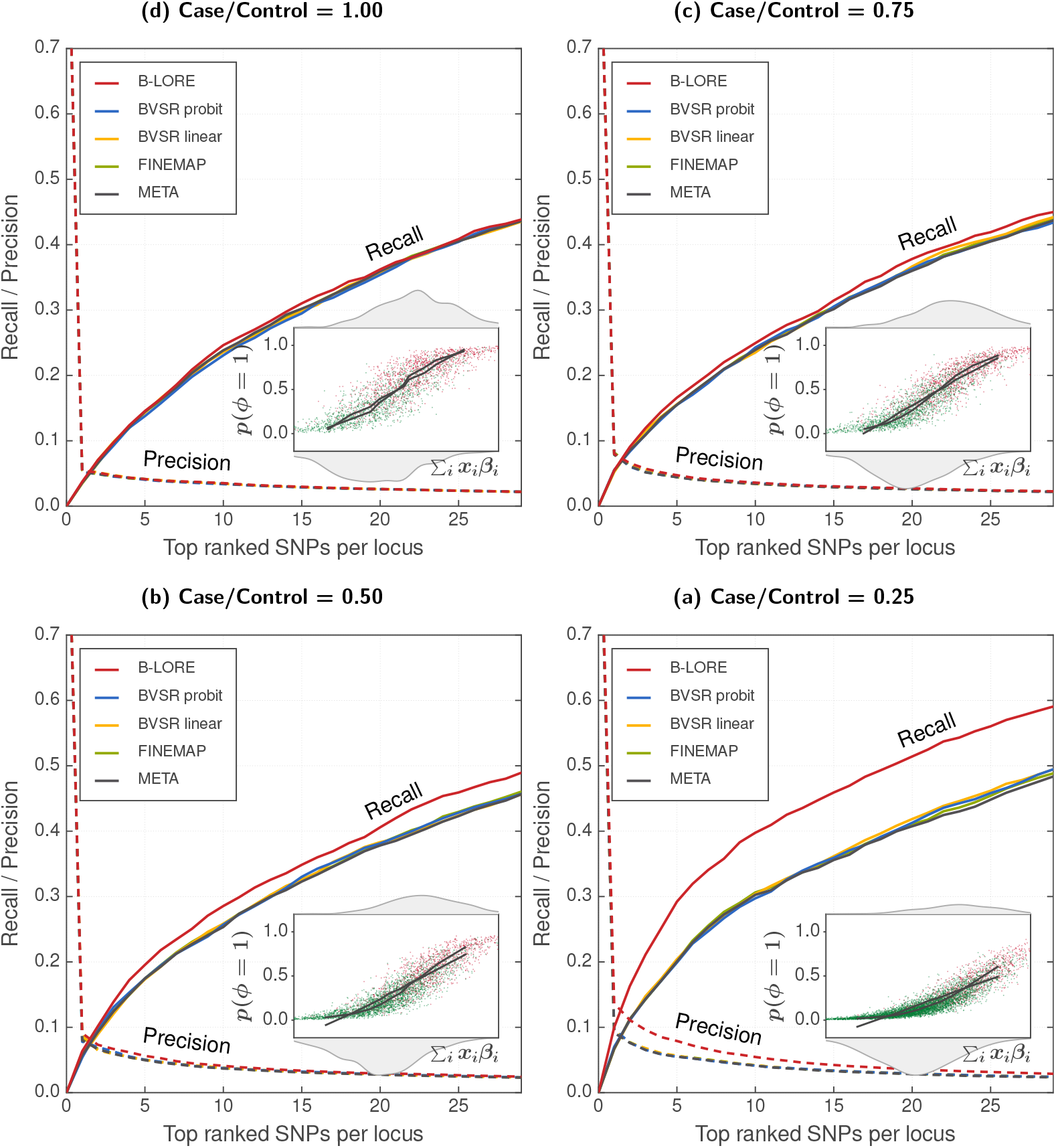
Multiple logistic regression improves power of GWAS with additional controls. We simulated phenotypes with varying case/control ratio – (a) 1625/1625, (b) 1625/3250, (c) 1625/4875 and (d) 1625/6500 respectively – using 100 loci of ~200 SNPs, as described in the main text. All simulations used 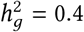. We compared the ranking of SNPs at each locus using recall (solid lines, left *y*-axis) and precision (dotted lines, right *y*-axis), which were averaged over 100 loci and 20 simulation replicates. All methods were run with a maximum of two causal SNPs per locus. Insets schematically compare logistic model with linear model (see Fig. 1 for details). B-LORE shows increasingly more recall over other methods with addition of more controls, *i.e.*, decreasing case/control ratio, because the logistic function becomes increasingly better than the linear function to model the data.

In both scenarios (a) and (b), B-LORE provides increasingly more recall over other methods as the case/control ratio decreases, confirming that the quasi-Laplace is more accurate than the existing linear approximations for the logistic model. Scenario (a) starts with 1625 cases and 1625 controls for 1:1 ratio. Due to less number of total samples, all methods have reduced power as compared to Fig. 1b. All methods provide almost similar recall for 1:1 case:control ratio. Including additional controls improves the recall of all methods but B-LORE gets the maximum benefit. When the number of controls is four times as much as the number of cases (case/control = 0.25), the logistic model has 33% better recall than the linear model for choosing the top 10 SNPs (and 45% better recall for choosing the top 5 SNPs).

In scenario (b), all methods have maximum recall when case/control = 1.0. With reducing number of cases, BVSR, FINEMAP and META lose power, while B-LORE remains virtually unaffected.

We also found that the BVSR probit model does not improve over the linear model. Even with a single locus, the probit model was expected to be better than the linear model in the extreme situation of case/control = 0.25. While we cannot rule out the effect of multiple loci, this result indicates that inefficient MCMC sampling is at least one of the reasons for the poorer performance of piMASS. By the same reasoning, the improved performance of B-LORE can be attributed to the logistic model although the effect of multiple loci cannot be ruled out.

### Effect of multiple loci

We then wanted to check the effect of using multiple loci on the improvement in B-LORE. Multiple loci together explain a higher proportion of heritability than each single locus. Newcombe *et al.* earlier showed that multiple-locus analysis is better at fine-mapping than many independent single-locus analyses [20]. Unfortunately, their method is limited to quantitative phenotypes and is not yet designed for binary phenotypes. Hence we could not include their method in our benchmark. Instead, we varied the number of loci in our analysis to directly check the effect of estimating causality from multiple loci.

We simulated phenotypes for the five cohorts with 25, 50, 75 and 100 loci – with 80, 40, 33 and 20 simulation replicates respectively. Each locus had ~200 SNPs. We used a fixed heritability of 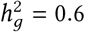 explained by these loci and 1:1 ratio of cases to controls.

In Fig. 3 we show the fine-mapping performance of the different methods in these simulations. The heritability per locus increases with decreasing number of loci. Hence we observe that the performance of all methods improves when the number of loci is reduced. However, B-LORE always provides higher recall than other methods.

**Fig 3.**
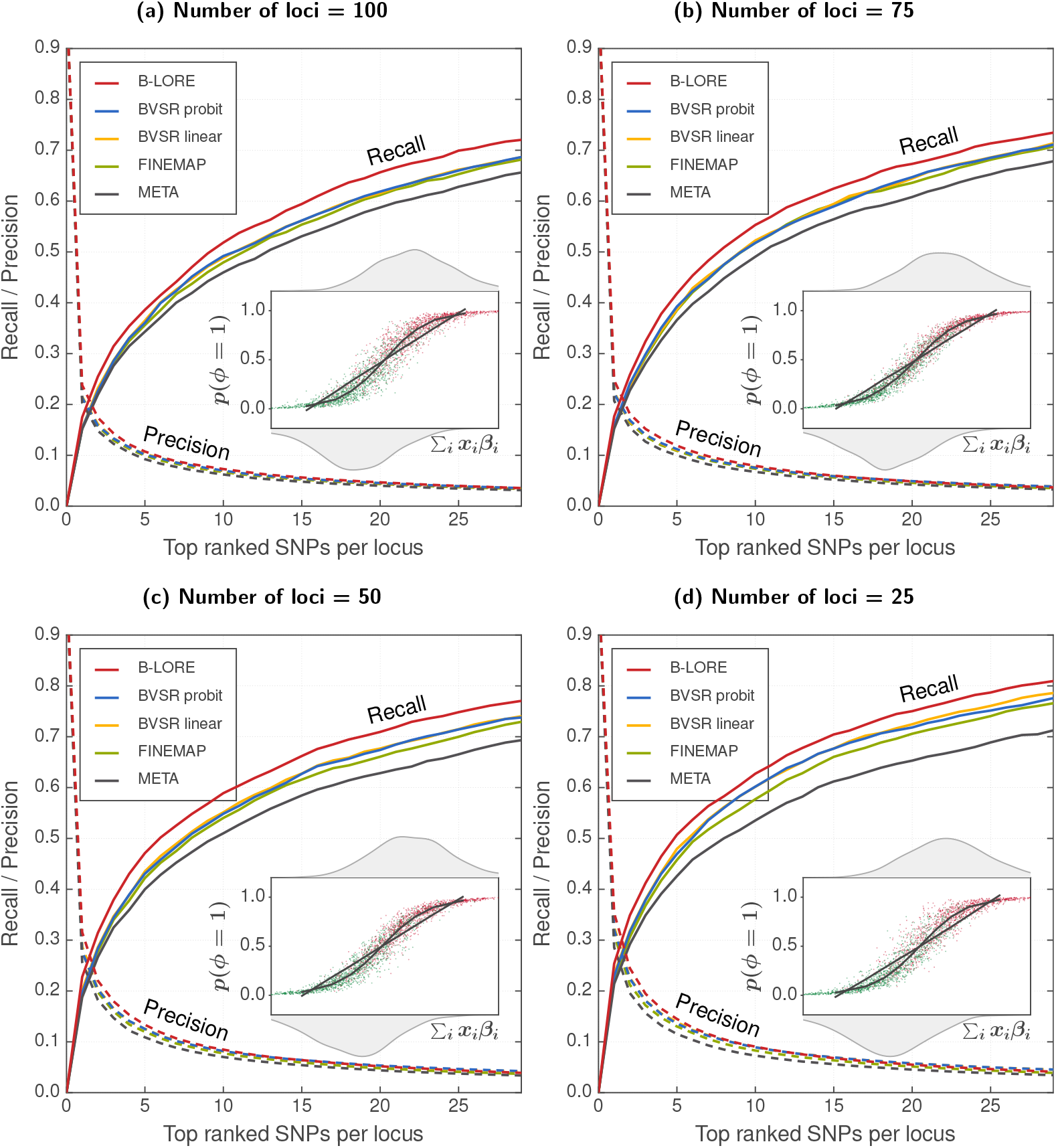
The advantage of B-LORE does not depend on the number of loci used for estimation. We simulated 13082 phenotypes using 25, 50, 75 or 100 loci of ~200 SNPs, as described in the main text. All simulations used 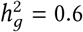. We compared the ranking of SNPs at each locus using recall (solid lines, left *y*-axis) and precision (dotted lines, right *y*-axis), which were averaged over the loci and the simulation replicates. All methods were run with a maximum of two causal SNPs per locus. Different panels show the results at different number of loci. Insets schematically compare logistic model with linear model in one simulation (see Fig. 1 for details). The heritability per locus increases when the number of loci is reduced. Multiple regression becomes increasingly better than single SNP analysis, but the advantage of B-LORE over other multiple regression methods does not change with the number of loci. Note also that the comparison between logistic model and the linear model in the insets does not change with the number of loci.

We further wanted to decouple the effect of multiple loci on calculating the summary statistics and meta-analysis of B-LORE. For this, we assumed that the phenotype is simulated from 100 loci as usual, and asked the following two questions. First, what happens if the B-LORE summary statistics are calculated from a subset of these 100 loci? This would mean that the heritability *visible* to B-LORE (*i.e.*, the heritability explained by the chosen subset of loci) is reduced, *e.g.* calculating summary statistics with 25 loci would correspond to a heritability of 0.15 when the total 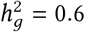. Hence the results (S8 Figure) are similar to Fig. 1. Second, what happens if the summary statistics are calculated from 100 loci for each cohort but the meta-analysis uses only a subset of them? In S9 Figure, we show that the ranking in each locus does not depend on the number of loci in the subset, indicating that the number of loci does not impact the estimation of hyperparameters, as long as we have enough number of causal SNPs (~ 20) in the data for estimating the hyperparameters. Therefore, we can conclude that our method benefits from the greater proportion of heritability explained by the multiple loci.

### Number of causal SNPs allowed

Having established that B-LORE improves the power of case-control GWAS over existing methods, we next explored the effect of allowing different numbers of causal SNPs ║**c**║_1_ in the model of B-LORE. Different loci are expected to have different number of true causal SNPs in the data. Current implementation of B-LORE allows only one common input ||c| _1_ for all loci.

There is no way to know the “ground truth”, *i.e.*, the distribution of true causal SNPs in the different loci of real GWAS. Here, we use different hypothetical distributions of true causal SNPs to simulate the phenotype and check the effect of ║**c**║_1_ on the ranking performance of B-LORE (Fig. 4). We plot the hypothetical distribution of true causal SNPs used for each simulation in the insets of each panel respectively. In general, higher c improves the ranking power of B-LORE, but does not significantly impact FINEMAP. Of course, the impact depends on the “ground truth”. For most practical cases, when there are few or no regions with > 5 truecausal SNPs, ║**c**║_1_ = 3 suffices for B-LORE.

**Fig 4.**
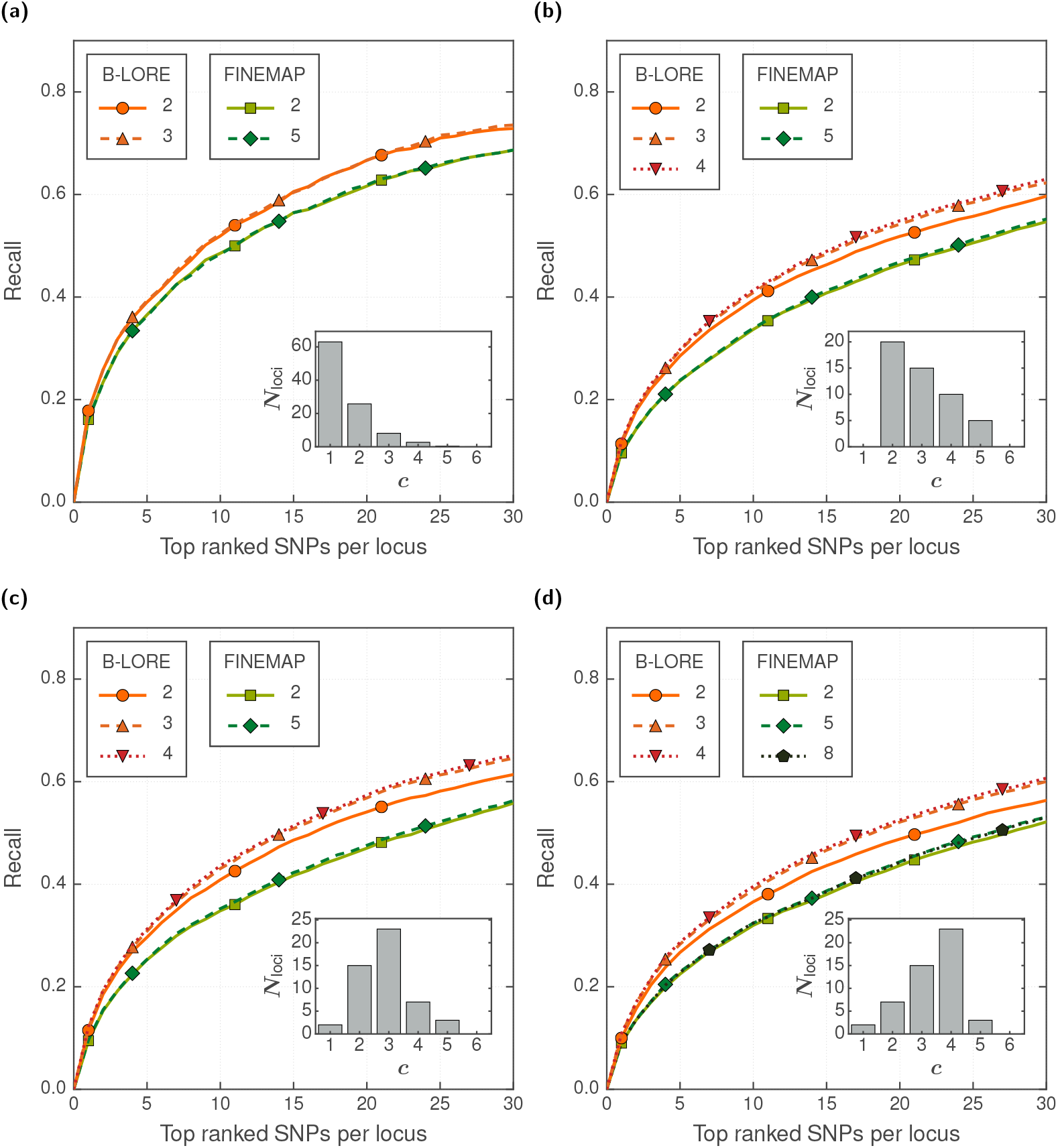
Effect of number of causal SNPs in B-LORE fine-mapping accuracy. We simulated 13082 phenotypes using 100 loci of ~200 SNPs, as described in the main text. Different panels show the results using different hypothetical distributions of true causal SNPs in each simulation (see insets, the distributions were generated *ad hoc*). All simulations used 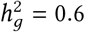. We compared the ranking of SNPs at each locus by B-LORE and FINEMAP using recall (solid lines, left *y*-axis) averaged over the loci and the simulation replicates. Both methods were run with different number of causal SNPs allowed in the model (║**c**║_1_ see legends). FINEMAP was run on each locus separately and B-LORE was run on all loci together. For each method, we stopped increasing ||c|| _1_ if the recall did not improve. The symbols are merely visual guides to distinguish between the different methods.

Earlier studies showed stronger influence of ║**c**║_1_ on fine-mapping accuracy (*e.g.* see Fig. 5 of [10]). However, those simulations used replicates of a single locus with fixed number of true causal SNPs. In contrast, we used many loci with a distribution of true causal SNPs because we expect that future applications of B-LORE would involve analyses over all loci found in a GWAS.

### Computational efficiency

In S6 Figure, we show the average time and memory required for calculating the summary statistics of B-LORE in different realistic situations. For instance, computing the summary statistics of 40000 SNPs spread over 200 loci in a cohort of 10000 individuals requires around 2 hours and ~22 GB of memory on an Intel Xeon E5-2670 v2 processor with 8 cores.

For B-LORE meta-analysis, we compare the average time and memory requirements of B-LORE with other multiple regression variable selection meta-analysis methods (CAVIARBF and FINEMAP) in S7 Figure. B-LORE has a speed comparable to CAVIARBF, although, unlike CAVIARBF, B-LORE optimizes the hyperparameters.

B-LORE is significantly slower than FINEMAP, and requires much more memory than FINEMAP. This is expected because B-LORE performs exhaustive search over the causality configurations, while FINEMAP uses a much faster shotgun stochastic search. The speed improvement by FINEMAP was a major technical breakthrough in multiple regression. We currently focus on better performance and only use a naive branch-and-bound algorithm (S1 File) to reduce the search space. Note that unlike FINEMAP, B-LORE learns *σ* from the data.

Like other multiple regression methods, B-LORE is not designed for genome-wide analysis. It could be used to finemap each locus as well as re-rank the loci according to their probability of being causal (see Inference). Considering the speed and memory requirements (S6 Figure and S7 Figure), we showed that with modest computing facilities, it is possible to apply B-LORE even on a total of 40000 SNPs, distributed evenly over 200 loci. The total number of SNPs and the total number of loci can be increased with more computing power, which is commonly available. There has to be a balance between the number of SNPs (*I_l_*) within a locus and the number of causal SNPs (║**c**║_1_) used by the model because the exhaustive search over the SNPs creates combinatorially increasing causal configurations depending on *I_l_* and ║**c**║_1_ For example, *I_l_* can be up to a few thousands with ║**c**║_1_ = 2, but *I_l_* can only be up to a few hundreds with ║**c**║_1_ = 5. Different loci might have different number of causal SNPs, and we showed that ║c║ _1_ = 3 is sufficient for a wide variety of distributions (Fig. 4).

### Calibration of posterior inclusion probabilities (PIPs)

Finally, we note that the PIPs obtained from B-LORE are well-calibrated (S5 Figure). Following the method proposed by Guan *et al.* [5], we assessed that the PIPs obtained from B-LORE roughly correspond to the marginal precision. On null data, B-LORE does not show any spurious association (S3 Figure).

## Discussion

Since its introduction in 2011, BVSR has remained the *de facto* method of choice for achieving the maximum power in GWAS multiple regression [5], but was limited by the requirement of individual genotype data. Several fine-mapping methods [9, 10, 30] introduced novel strategies to extend the scope of multiple regression analysis using only summary statistics from individual studies. However, none of these methods achieved more power [9,30] than BVSR unless additional external information, such as ENCODE data or multiple traits, was used [6, 7, 31].

In the present work, we introduce B-LORE for performing multiple logistic regression on binary phenotypes, and show that it improves the power of case-control GWAS over both BVSR probit and linear models. In fact, the improvement achieved by B-LORE over BVSR is more significant than the improvement achieved by BVSR over single-SNP analyses (Fig. 1). The key innovation is the quasi-Laplace approximation, which allows us to accurately compute posterior probabilities for SNP causality and to learn hyperparameters for the multiple logistic regression model.

Multiple logistic regression has received very little attention in GWAS. There were technical difficulties in the MCMC sampling of binary phenotypes for the BVSR probit model. The fine-mapping methods approximated the logistic likelihood with a Gaussian, which is essentially equivalent to using a scaled linear model. This linear approximation becomes inaccurate when moving out of the inflection point of the logistic function or when the spread of disease risk among patients is large.

We can get some intuition about this advantage of the logistic over the linear model from the partial derivatives of their log likelihoods with respect to the effect sizes, ***β***,

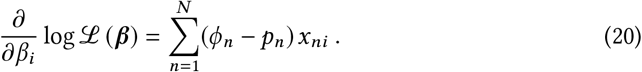

Here, *p_n_* is given by Eq. (1) for the logistic model and by *p_n_* = ***β***^Τ^**x**_*n*_ for the linear one. The terms (*φ_n_* – *p_n_*) can be interpreted as weights with which diseased patients (*φ* = 1) with minor alleles at SNP *i* (*x_ni_* > 0) “vote” *β_i_* up and healthy patients (*φ* = 0) with *x_ni_* > 0 “vote” *β_i_* down. The strongest contributions are made by healthy patients with high predicted disease risk pn and by diseased patients with low predicted risk, while the smallest contributions are made by healthy patients with low predicted risk and diseased patients with high predicted risk. As long as pn is near 0.5, the weights for both models are near 0.5 and they will give very similar results. But as predictability grows or the bias changes from 0.5 due to case/control ratios different from 1, the *p_n_* for the logistic model will become quite different from those of the linear model and the linear model will become inaccurate. In extreme cases when ***β***^Τ^**x**_*n*_ gets smaller than 0 or larger than 1, patients will even “vote” with the wrong sign!

When applying logistic regression to a single SNP or when applying multiple logistic regression only on a single locus at a time, the predicted risk differences are small, the values of *p_n_* lie very near to 0.5, and the linear approximation holds well. However, when predicted risks lie in the non-linear regime of the logistic curve, B-LORE outperforms – by a clear margin – methods based on a linear regression model and those based on a linear approximation of the logistic regression model. We gave two examples:

First, when applying multiple logistic regression to all risk loci, heritability adds up over the many loci, the predicted risks *p_n_* scatter farther from 0.5 and the logistic model performs significantly better than the linear model (Fig. 1, S8 Figure).

Second, when the case-control ratio r differs significantly from 1, the predicted risk of patients scatters around 1/(1 + *r*), which will differ significantly from 0.5. In this regime, B-LORE strongly outperformed the multiple linear regression methods (Fig. 2). The failure of the linear model in the nonlinear regime could explain the reason why GWAS have been rarely analyzed with unbalanced case-control ratios. B-LORE could change that. It should be possible to discover many new loci and SNPs associated with complex diseases simply by using additional controls to existing GWAS, *e.g.* from one of the medical biobanks springing up across the world [32].

Third, predicting the disease risk from known covariates should significantly improve power of detecting causal SNPs, because the more accurate estimation of *p_n_* would improve the weighting of patients. It is well known that age and sex alone can predict disease risk more accurately than genotype for most common diseases (e.g. cardiovascular diseases [33,34]). Various other measurable covariates could further improve risk prediction. Using covariates to predict disease risk from external data sources [35] (and *not* from the case-control study itself, as this estimate would suffer from strong selection bias) could therefore greatly improve power in combination with our logistic multiple regression approach.

The flexibility of the Bayesian approach makes it easy to integrate this and other external information in the future, such as functional genomics data tracks on DNA accessibility, transcription factor binding etc. [6,36–38] which can modulate the prior probability *π*_i_ for a SNP to be causal. We will explore more systematically how best to improve predictive performance of multiple logistic regression by adding these functional annotations.

The quasi-Laplace approximation also allows us to extend the analysis to multiple studies without using genotype data. Any method for GWAS meta-analysis that can improve the power has enormous leverage. It can be applied to pool millions of patients from multiple GWAS to help understand the origin of all common diseases in humans [2]. We hope that B-LORE will contribute to realizing this potential.

## Acknowledgments

The comments of the anonymous reviewers of an earlier version of the manuscript encouraged us to explore the validity and utility of the quasi-Laplace approximation. We thank them for their constructive feedback. We thank Franco L. Simonetti and Anubhav Kaphle for feedback on the manuscript, Salma Johrabi for feedback on the figures, Clovis Galiez, Gonzalo Parra and Christian Roth for helpful discussions. This work was supported by the German Federal Ministry of Education and Research (BMBF) within the framework of the e:Med research and funding concept (grants e:AtheroSysMed 01ZX1313A-2014).

## Supporting information

### S1 File. Additional Methods

#### S2 Figure. Distribution of the causal SNPs in our simulation

In each locus, we sampled the causal SNPs with a prior probability *π* and a constraint that at least one SNP must be picked. For our simulations, we choose *π* = 0.005. We show (a) the distribution of total number of causal SNPs used for each simulation, and (b) the distribution of causal SNPs in each locus averaged over all simulations.

#### S3 Figure. Performance of B-LORE on null data

We performed a simulation with randomly generated binary phenotype on 13082 samples across five populations, using 17218 SNPs distributed over 100 loci. We show (a) the log posterior probability of each locus being causal (log_10_ (Pr_causal_)), and (b) the posterior inclusion probability (PIP) for each SNP.

#### S4 Figure. Imbalance in case-control GWAS with fixed number of controls

We simulated phenotypes with varying case/control ratio – (a) 1625/6500, (b) 3250/6500, (c) 4875/6500 and (d) 6500/6500 respectively – using 100 loci of ~200 SNPs, as described in the main text. All simulations used 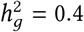. We compared the ranking of SNPs at each locus using recall (solid lines, left *y*-axis) and precision (dotted lines, right *y*-axis), which were averaged over 100 loci and 20 simulation replicates. All methods were run with a maximum of two causal SNPs per locus. Insets schematically compare logistic model with linear model (see Fig. 1 for details). B-LORE shows increasingly more recall over other methods with increasing imbalance, because the logistic function becomes increasingly better than the linear function to model the data.

#### S5 Figure. Calibration of the posterior inclusion probabilities from B-LORE

The SNPs were put into 10 bins of width 0.1 according to their posterior inclusion probabilities (PIPs). Each point on the plot represents a single bin, with the center of the PIP within that bin on the *x*-axis and the proportion of SNPs which were true positives in that bin on the *y*-axis. Vertical bars show ±2 standard errors of the proportions, assuming a binomial distribution. Panel (a) is the result of B-LORE using maximum of 2 causal SNPs and panel (b) using maximum of 3 causal SNPs.

#### S6 Figure. CPU time and memory requirement for calculating summary statistics with B-LORE

Panel (a) shows the processing time and panel (b) shows the maximum memory required for calculating B-LORE summary statistics with 50, 100, 150 and 200 loci. Each point corresponds to an average over 20 simulations, with the vertical bars representing ± standard errors. Different shapes and colors correspond to different sample sizes (*N* = 2000, 4000, 6000, 8000 and 10000) of GWA studies, as specified in the legend. The lines connect the results of the same sample size. Each locus contained 200 SNPs. All calculations were done on an Intel Xeon E5-2670 v2 processor with 8 cores.

#### S7 Figure. CPU time and memory requirement for meta-analysis with B-LORE

We compared the computational requirements of B-LORE with other fine-mapping methods in terms of (a) processing time and (b) maximum memory required. Along the *x*-axis, we vary the number of maximum allowed causal SNPs. For each point on the plot, we used an average over 20 simulations. Each simulation was a meta-analysis of 5 GWA studies with 40000 SNPs (distributed over 200 loci). All calculations were done on an Intel Xeon E5-2670 v2 processor with 8 cores. FINEMAP and CAVIARBF were allowed to use all the cores in parallel, by analyzing 25 loci in each core.

#### S8 Figure. Impact of number of loci on calculation of B-LORE summary statistics

We simulated 13082 phenotypes using 100 loci of ~200 SNPs, as described in the main text. All simulations used 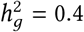. We then used only a subset (25, 50, 75 and 100) of these loci for further analysis. We compared the ranking of SNPs at each locus using recall (solid lines, left *y*-axis) and precision (dotted lines, right *y*-axis), which were averaged over the loci and the simulation replicates. All methods were run with a maximum of two causal SNPs per locus.

#### S9 Figure. Impact of number of loci on optimization of hyperparameters for B-LORE meta-analysis

We simulated 13082 phenotypes using 100 loci of ~200 SNPs, as described in the main text. All simulations used 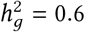. We calculated B-LORE summary statistics from all the 100 loci, but performed meta-analysis using only a subset (25, 50, 75 and 100 – shown in different panels) of these loci. Other methods were run on the same subset of loci. We compared the ranking ranking of SNPs at each locus using recall (solid lines, left *y*-axis) and precision (dotted lines, right *y*-axis), which were averaged over the loci and the simulation replicates. All methods were run with a maximum of two causal SNPs per locus.

